# Narrow, but not broad, spectrum resistance and disease reshape phyllosphere bacterial communities

**DOI:** 10.1101/2023.08.11.551834

**Authors:** Adam F. Bigott, Samuel F. Hutton, Gary Vallad, Rick Lankau, Jeri Barak

**Author notes:** Corresponding author: Jeri Barak.

## Abstract

The phyllosphere is a restrictive environment for microbes, resulting in microbial communities typically dominated by select taxa with specific adaptations for success in this niche. However, biotic stress, especially from plant disease, could disrupt this environment in ways that alter the resulting phyllosphere community, with potential consequences for plant and human health. Additionally, plant disease resistance, through both broad (pattern-triggered immunity) and specific (effector-triggered immunity) resistance, could affect non-pathogenic communities directly or indirectly via effects on disease progression. Here, we tested how transgenic ETI and PTI resistance genes affected the phyllosphere communities of tomato plants in the face of infection by *Xanthomonas perforans* and the resulting bacterial spot disease. We found that the expression of the *Bs2* transgene (ETI) had major effects on phyllosphere communities, while the EFR (PTI) transgene did not. The effect of the *Bs2* resistance gene could be largely attributed to the change in disease symptoms. Diseased leaves harbored reduced bacterial diversity and reductions in major phyllosphere inhabitants (e.g. *Sphinogmonas* and *Methylobacterium*), while a limited number of bacterial genera showed increased relative abundance on diseased leaves. These results suggest that phyllosphere communities are sensitive to the direct and indirect effects of plant disease and resistance, and the consequences of these shifts for plant and human health deserve further investigation.

## Introduction

The phyllosphere, or above-ground portion of a plant, is a restrictive ecological niche for microbes. As such, phyllosphere microbial communities are typically dominated by a restricted set of taxa with specific adaptations to this challenging environment. However, phyllosphere environments are also dynamic, subjecting microbial communities to fluctuating physical conditions as leaves age, are buffeted by weather events, or are attacked by herbivores and pathogens. While a relatively consistent view of the“healthy” phyllosphere has emerged from surveys across disparate plant species (Delmotte et al. 2009; Knief et al. 2012; Allard et al. 2016), how the phyllosphere microbial community changes in response to biotic stress on leaves remains unclear. This may be especially important as phyllosphere communities can contain rare and infrequent members that nonetheless have important consequences, including potential human pathogens, and whose populations may benefit from alterations to the phyllosphere environment.

Leaves are covered in waxy cuticles that limit the availability of moisture in the phylloplane, the leaf surface (Beattie 2011). Ultraviolet stress on the phylloplane leads to diversification of the associated bacterial communities favoring those with pigmentation and/or copious extra polymeric substance production (Kadivar and Stapleton 2003). High light and low humidity decrease bacterial phyllosphere populations (Wilson et al. 1999). Further, the availability of carbon sources in the phylloplane is limited and aggregated (Leveau and Lindow 2001). The aggregation of available nutrients results in few sites on the leaf surface where bacteria can replicate and reward motility and chemotaxis towards a more hospitable oasis such as glandular trichomes or host cellular junctions (Remus-Emsermann et al. 2012).

Even in this restrictive environment, bacteria are the most abundant members of the phyllosphere microbiome and can reach population densities as high as 10^7^ cells per cm^2^ or 10^8^ cells per gram of leaf tissue (Lindow and Brandl 2003). A small number of taxa, including *Sphingomonas*, *Methylobacterium*, and *Pseudomonas*, have been shown to be common members of the phyllosphere bacterial community across plant species, including tomato, rice, *Arabidopsis*, clover, and soybean (Delmotte et al. 2009; Knief et al. 2012; Allard et al. 2016). Carbon availability limits epiphytic bacterial populations (Wilson and Lindow 1995); these taxa have made different adaptations primarily in carbon acquisition and/or utilization to succeed in the phyllosphere. A combined metagenomic and metaproteomic analysis assessing bacterial communities on *Arabidopsis*, clover, and soybean described a large number of TonB dependent transporters assigned to *Sphingomonas* for putative uptake and utilization of a diverse array of carbon substrates (Delmotte et al. 2009), suggesting that the success of *Sphingomonas* in the phyllosphere is in part due to scavenging various substrates present at low amounts. Delmotte et al. (2009) also found *Methylobacterium* produced much higher levels of the proteins required for methylotrophy as compared to previously reported values when grown in culture.

*Pseudomonas* differs from the other two cosmopolitan phyllosphere genera in that it contains both non-pathogenic and pathogenic members. In order to deal with waxy, hydrophobic leaf surfaces, *Pseudomonas* produces biosurfactants to increase the wettability of the leaf for enhanced diffusion of nutrients across the waxy cuticle and/or aid in motility to favorable colonization sites (Burch et al. 2014). As a plant pathogen, *Pseudomonas syringae* has evolved different strategies for proliferation in the phyllosphere such as the production of coronatine to prevent stomatal closure for apoplastic entry (Zheng et al. 2012), the first step in the infection cycle, and in some cases, conversion of the apoplastic space to an aqueous environment. Accessing the apoplast alleviates stresses associated with the phylloplane and may have more available free water and nutrients.

Yet as bacteria interact with plant cells in apoplastic spaces, they encounter an additional challenge to survival, the plant immune system. Bacterial pathogens often suppress host defenses by disruption of immune signaling and other metabolic functions. During infection of tomato, *P*. *syringae* uses a type 3 (T3) effector (HopM1) to induce establishment of an aqueous apoplast, presumably increasing nutrient availability and potentially dampening pathogen associated molecular patterns (PAMP)-triggered immune (PTI) responses (Xin et al. 2016). Phyllosphere modification is not unique to pathogenic pseudomonads. On infection of tomato, *Xanthomonas gardneri* reprograms host pectate lyases by secretion of AvrHah1, a transcription activator-like T3 effector that also alters the apoplast to an aqueous environment (Schwartz et al. 2017). Furthermore, *X. translucens* infection of wheat leads to transcriptional reprogramming that converts infected leaves from carbohydrate sources to sinks (Garcia-Seco et al. 2017). In addition to modification of the phyllosphere, *Xanthomonas* effectors also contribute directly to pathogen fitness. Following the deletion of *avrBS2*, inoculation of the susceptible pepper cultivar Early Calwonder with *X. vesicatoria* populations were 100 times smaller after four days than their wildtype counterparts (Kearney and Staskawicz 1990).

Changes to the phyllosphere niche from infection benefit the pathogen and can alter colonization success for other bacteria. *Xanthomonas*-infection of tomato increases *Salmonella enterica* survival in the phyllosphere and results, in some cases, in bacterial replication (Potnis et al. 2014; Potnis et al. 2015). However, on hosts resistant to *Xanthomonas*, the human pathogen population decreased rapidly. Without changes to the phyllosphere caused by infection, *S. enterica* has limited success, fails to replicate, and immigrant populations decline over time (Islam et al. 2004a; Islam et al. 2004b; Potnis et al. 2014). How the bacterial community broadly responds to niche alterations due to disease or the triggering of host resistance by pathogens remains largely unknown.

Our aim in this study is to analyze how the bacterial phyllosphere community responds to host infection and disease resistance. We used a well-characterized pathosystem of *Xanthomonas perforans* and tomato. *X. perforans* causes bacterial spot of tomato and pepper, leading to significant changes to the phyllosphere environment, including establishment of an aqueous apoplast. There is little resistance in tomato to bacterial spot, but pepper, a close relative to tomato and fellow host to bacterial spot, has several resistance genes, including *Bs2* (Tai et al. 1999). Resistance to *avrBs2-*carrying strains of *Xanthomonas* spp. in pepper is conferred in a gene for gene interaction with the nucleotide binding leucine rich repeat receptor *Bs2* (Minsavage et al. 1990). Transgenic expression of *Bs2* from pepper into tomato was shown to both reduce disease severity and increase yield in field trials (Lacombe et al. 2010; Horvath et al. 2012). Plants can also initiate a broad-spectrum resistance to bacterial pathogens when Receptor-Like Kinases on host cells are activated by binding to broadly conserved microbial epitopes, such as flagellin or elongation factor tu (Jones and Dangl 2006; Boller and Felix 2009). Tomato lines expressing the *Arabidopsis thaliana* gene *EFR*, a PAMP receptor recognizing elongation factor tu, have been developed to resist multiple bacterial pathogens simultaneously. *EFR-*expressing plants are resistant to diseases caused by *P. syringae*, *Agrobacterium tumafaciens*, and *Ralstonia solanacearum* (Lacombe et al. 2010; Kunwar et al. 2018). While transgenic PTI precludes disease development, the effects on non-pathogenic microbes is poor understood. Phyllosphere community assembly is partially shaped by early and influential colonizers (Debray et al. 2023). Disruption of this process by transgenic *EFR* could lead to distinct and unforeseen differences in phyllosphere community composition.

In this study, we characterized the phyllosphere bacterial community of tomato using 16S metabarcoding. We used isogenic lines of tomato varying in the presence or absence of both a narrow– and a broad-spectrum resistance gene, grown using commercial production standards under high disease pressure. To correlate bacterial community composition with disease, we also measured *Xanthomonas* disease severity and fruit yield. Using this information, we examined the importance of the pathogen and/or host resistance in shaping bacterial community structure. We calculated the relative abundance of bacterial community members to examine how cosmopolitan genera, *Sphingomonas*, *Methylobacterium*, and *Pseudomonas*, were affected by host genotype and thus, disease.

## Materials and Methods

*Sampling and DNA Processing:* The tomato plants used in these experiments were grown at and managed by the University of Florida Gulf Coast Research & Education Center in Balm, Florida. Fields were prepared according to standard commercial field production with fumigated (Pic-Clor 60, 336 kg/ha) raised beds (81.3 cm wide at the base, 71.1 cm wide at the crown and 20.3 cm in height) in groups of three at 91.4 m lengths, spaced on 1.5 m row spacing, and covered with either a white (fall) or black (spring) virtually impermeable film. Plots were arranged in a randomized complete block design with each treatment represented by four blocks of 10 plants per plot. The experimental design consisted of four plant genotypes (Fla. 8000, Fla. 8000 *BS2*, Fla. 8000 *BS2 EFR*, and Fla. 8000 *EFR*) crossed with an *Xanthomonas perforans* inoculation treatment. Inoculated treatment plots were inoculated with a cocktail of *Xanthomonas perforans* race T4 strains: GEV904, GEV917, GEV1001, and GEV1063. Inoculations and mock inoculations were performed two weeks after transplanting 4-week old seedlings at 45.7 cm spacing. Non-inoculated plots were mock inoculated with an equal volume of water. Leaf samples were collected at the end of the season and 2 mature leaves from the upper canopy of each of the 10 plants in a plot were binned for a single sample. Samples were shipped on ice and processed within 24 hours of collection. The experiment was repeated during three separate field seasons: fall 2017, spring 2018, and fall 2018.

Leaf fresh weights were recorded prior to leaf wash extraction. Leaves were double bagged in ziplock bags with 600 ml of water and 12 μl Hi-Wett (Loveland Products Inx., Greeley, CO). Leaves were washed in bags by vortexing on high for 30 seconds and then sonicated for 30 seconds using a Branson 5800 sonicator in two successive cycles. Leaf wash was then transferred to a Thermo Scientific Nalgene Analytical Test Filter Funnel and filtered with a vacuum pump. Filters containing leaf extract were cut into quarters and stored in 2 ml microcentrifuge tubes at –80C.

DNA extraction from the leaf extract filter paper was performed using the Omega Bio-Tek E.Z.N.A. Plant DNA Kit following the manufacturer’s instructions. Extracted DNA was used as a template for an initial PCR amplification using 16S primer pair for the V5-V6 region (799F and 1115R) that do not amplify the chloroplast 16S sequences (Laforest-Lapointe et al. 2017; Redford et al. 2010). As previously described in Xue et al. 2019, a second PCR was performed while adding one of 8 variable length barcodes (5-8 bp) to each sample and space for the Nextera primer to bind. The external fusion PCR primers for the second PCR contained a 14 bp overlap with the end of the internal primer as well as one of two options: an 8-bp i7 index and P7 flow cell adapter sequence or a 7-bp spacer i5 index, and P5 flow cell adapter sequence.

The first PCR reaction used a total volume of 10 µL. Reactions used 0.2 µL of hot-start high fidelity Clonetech PrimeSTAR GLX polymerase (Takara Bio, Kusatusu, Japan), 2 µL of 5X buffer, 0.8 µL of 10 nM dNTPs, 0.25 µL of 10 nM forward and reverse primer, 0.7 µg T4 gene 32 protein, and 10 ng of template DNA. The Thermocycler protocol was as follows: a hot start at 98°C for 5 minutes, 35 cycles of denaturing at 98°C for 45 seconds, annealing at 50°C for 45 sec., extension at 68°C for 1 min., and a final 15 min. extension at 68°C.

The second PCR reaction used a total volume of 25 µL. Reactions contained 0.5 µL of hot-start high fidelity Clonetech PrimeSTAR GLX polymerase (Takara Bio, Kusatusu, Japan), 5 µL of 5X buffer, 1 µL of each 10 nM primer, and 1 µL of product from the previous PCR as template. The Thermocycler protocol was as follows: a hot start at 98°C for 5 minutes, 15 cycles of denaturing at 98°C for 30 seconds, annealing at 60°C for 0:45, extension at 68°C for 1:00, and a final extension of 10 minutes at 68°C.

Successful amplification at each step was visually confirmed on a UV-transilluminator following gel electrophoresis of the product in 1% agarose and staining with gel red. Final amplification products were purified with the Omega BioTek E-Z 96 Cycle Pure kit and quantified using a Qubit 2.0 fluorometer with the Qubit dsDNA HS assay (Thermo Scientific, Grand Island, NY). DNA was pooled at equal concentration and sent for Illumina MiSeq sequencing at the University of Wisconsin Biotechnology Center (Madison, WI).

*Processing of Sequencing Data:* External barcodes were demultiplexed by the Wisconsin Biotech Center. Each set of matching external barcodes contained multiple samples distinguished by the third barcode sequence between the Nextera adapter and 16S primer sequence. This second demultiplexing was performed in Qiime2 (Boylen et al. 2018) using the cut-adapt plugin from the demux-paired function.

Denoising, replication, and chimera filtering were all performed in Qiime2 using the dada2 plugin’s “denoise-paired” option using default settings. Taxonomic assignment was performed using naïve Bayesian classification. The Greengenes database was used as a reference database for taxonomic identities (McDonald et al. 2012). Following assignment, a dataset for downstream analysis was created by combining amplicon sequence variants identified to the same genus. Sequences that could not be identified to a given taxonomic resolution were lumped into a single unclassified taxon with terminal identification at the next highest taxonomic rank.

Illumina sequencing of 16S amplicons are archived in the sequence read archive under BioProject accession number PRJNA604617. In order to address a potential bias in community diversity rooted in differential sequencing depth between plots, samples were assessed using rarefaction curves of the number of genera (Fig. S1). In order to retain as many samples as possible, samples were rarefied to an even depth of 10,000 sequences. Each season of collected data contained 32 samples. Of the 96 total samples collected, 84 were retained at this sampling depth. The results of rarefaction show that not all samples exhibit a plateau in the number of genera present at the final sampling depth, suggesting that some samples may be underrepresented in more rare taxa. Across all three seasons of data, a total of 691 distinct genera were detected.

*Yield Analysis:* Tomatoes were harvested by making four separate picks between 10 to 12 weeks after transplanting. Fruit were graded and sorted into the following categories according to USDA grading standards: Culls, Small, Medium, Large, and Extra-large. Cull and small fruits were not incorporated into marketable yield estimates used in this analysis. Since the number of picks made to assess yield varied from season to season, the total yield for each season was analyzed independently.

*Bacterial Spot Disease Rating:* Individual plants within each plot were assessed for bacterial spot using the Horsfall-Barratt scale (Horsfall and Barratt 1945) and transformed to the midpoint value of each interval prior to statistical analysis (Redman et al. 1964).

*Statistical Analyses:* All statistical analysis was performed in R (version 4.1.0). Transformed averages of disease severity per plot, average fruit yield, and alpha diversity metrics were analyzed by performing a mixed effects linear model using the lmer and lmerTest packages in R followed by post-hoc Tukey’s HSD test in R using a value of p < 0.05. Mixed models included block nested within season-by-year as random effects. The assumptions of normality were assessed by visual inspection of residual value histograms to confirm a normal distribution of model residuals.

Permutational Multiple Analysis of Variance (PERMANOVA) was used to compare bacterial community composition among genotypes at the genus level. Bacterial community structure comparisons were visualized using distance-based redundancy analysis (db-RDA) of the Bray-Curtis dissimilarity matrix to visualize differences in community structure among bacterial phyllosphere communities.

We expected *Xanthomonas* to be a highly abundant taxa in our dataset, and to respond strongly to the presence of the BS2 gene. To determine whether any effects seen on phylosphere community structure stemmed solely from changes to the Xanthomonas population, we created a subsetted dataset that excluded Xanthomonas and rerarefied to a common sequence depth of 2,992 sequences. We then used the same PERMANOVA models as above to quantify the effect of the season, the *BS2* gene, and the EFR gene and their interactions on the composition of the non-Xanthomonas community.

The presence of the *BS2* gene could affect phyllosphere community composition due to the direct effects of *BS2* gene expression or due to indirect effects mediated through the Xanthomonas population and consequent disease symptoms. To disentangle these possibilities, we ran PERMANOVA models that included both the *BS2* gene treatment and the quantitative assessment of bacterial spot disease severity. We ran this model in two ways: once with the *BS2* gene treatment entered first in the model, and once with the disease severity term entered first. By comparing the resulting R^2^ and P values associated with these terms when entered first or second in the model we can assess how much variation in bacterial community composition can be attributed uniquely to one or the other term, and how much of their effects cannot be distinguished. As before, we did this on the entire community dataset, as well as a subset that excluded Xanthomonas.

Individual taxonomic relationships were assessed using rarefied metabarcoding data. Taxa were tested whether they differed in their relative abundance based on the presence absence of a transgene by Wilcoxon Rank Sum test to determine if their abundance differed in *Bs2* and non-*Bs2* plants as well *EFR* and non-*EFR* plants. Wilcoxon Rank Sum P values were corrected using False Discovery Rate (FDR) based on the number of applicable rank sum comparisons.

In order to help determine the certainty of the assigned taxonomic identity of OTUs for *Salmonella*, sequences of closely related taxa were obtained from Greengenes database (https://greengenes.secondgenome.com/) and aligned in Mega 7.0. Aligned sequences were then used to construct a maximum likelihood phylogeny. Identification of an OTU to genus was contingent on it being placed within a monophyletic clade of that genus.

## Results

*Inoculated and non-inoculated treatments did not differ with respect to field and sequencing data*. An initial analysis was conducted to test whether *X. perforans* inoculated and non-inoculated plots differed. ANOVA of yield, disease rating, proportion of *Xanthomonas* in the metabarcoding, and PERMANOVA testing bacterial community structure were not affected by the inoculation treatment (p > 0.1). Therefore, inoculated and non-inoculated treatment levels were combined in subsequent analyses.

*Transgenic resistance increased yield and lowered disease ratings*. Bacterial spot disease severity differed based on the presence of *Bs2* and *EFR* (Table 1). In all three seasons sampled, *Bs2* expressing plants were rated as having less than 1% disease severity. Disease severity in *EFR* expressing plants ranged from as low as 12% in the fall of 2017 to 47% in spring 2018 whereas WT Fla. 8000 plants ranged from 24% to 68%. WT plants and plants only expressing *EFR* had significantly higher disease in the spring of 2018 than either fall season (Fig. 1; Tukey’s HSD p < 0.05). Plants expressing either or both *Bs2* and *EFR* displayed reduced symptoms compared to WT plants and no differences in symptoms were observed between *Bs2* and *Bs2*/*EFR* expressing plants (Table S1).

**Figure 1.**
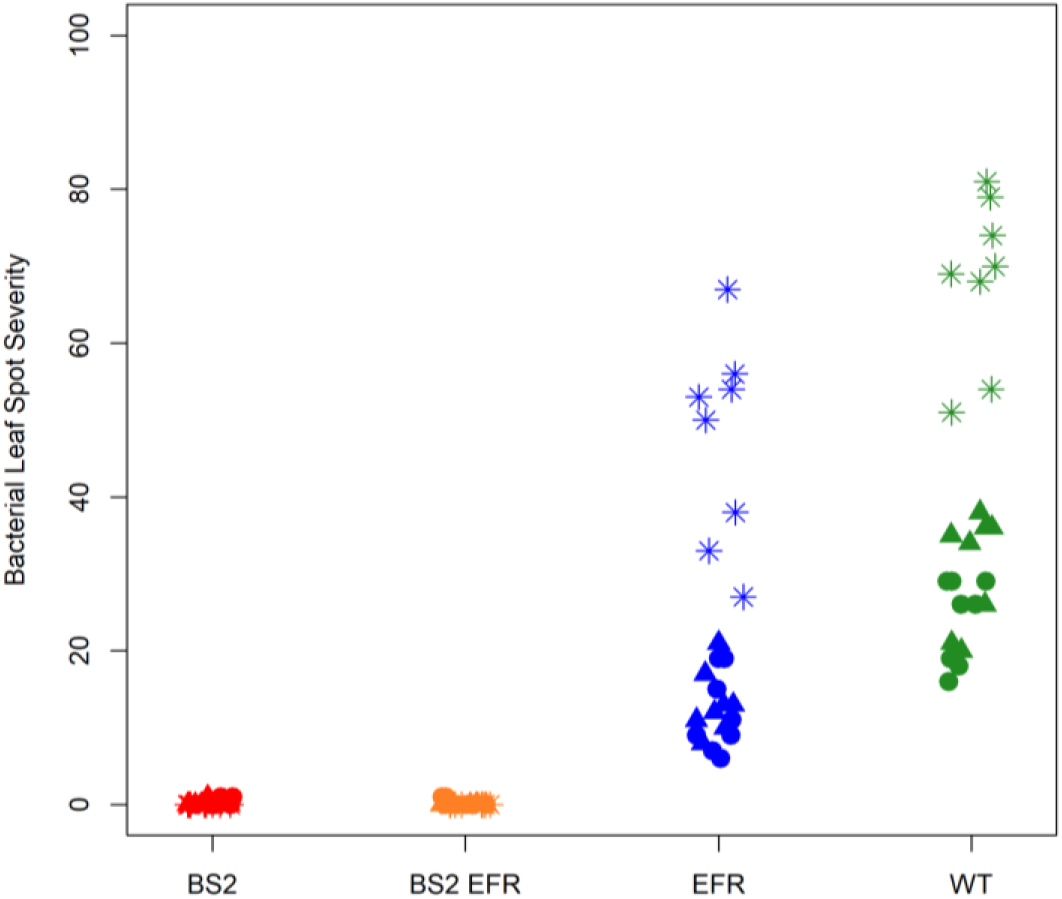
Little to no bacterial leaf spot was found on *Bs2* expressing plants and more disease occurred during the spring crop. Bacterial leaf spot severity average per plot. Color designates genotype (green, WT; blue, *EFR*; yellow *Bs2*/*EFR*; red, *Bs2*) and symbol sampling date (circle, fall 2017; asterisk, spring 2018; triangle, fall 2018). A one-way, analysis of variance (ANOVA) was used to assess transformed averages of disease severity per plot and average fruit yield, Tukey’s HSD p < 0.05.

**Table 1:** Little to no bacterial leaf spot was found on *Bs2* expressing plants and more disease occurred during the spring crop. A) The Horsfall-Barrat scale was used to estimate the severity of disease based on lesion size, number, and damage from disease progression to the leaf understory. Scores of individual plants were transformed to the midpoint of that interval before being averaged to produce disease severity values per plot. Relative yield was calculated for each plot based on the average yield across all plots per season.

| Year | Season | Genotype | Disease Severity | Fruit Yield (kg) |
| --- | --- | --- | --- | --- |
| 2017 | Fall | WT | 24.0 A | 18.1 B |
|  |  | *EFR* | 11.9 B | 18.5 B |
|  |  | *Bs2* | 0.3 C | 42.6 A |
|  |  | *Bs2/EFR* | 0.3 C | 40.4 A |
| 2018 | Spring | WT | 68.3 a | 44.5 ab |
|  |  | *EFR* | 47.3 b | 41.7 b |
|  |  | *Bs2* | 0.0 c | 54.6 a |
|  |  | *Bs2/EFR* | 0.0 c | 55.8 a |
| 2018 | Fall | WT | 30.8 $\alpha$ | 13.8 $\alpha\beta$ |
| | | *EFR* | 13.1 $\beta$ | 8.3 $\beta$ |
| | | *Bs2* | 0.1 $\chi$ | 20.1 $\alpha$ |
| | | *Bs2/EFR* | 0.0 $\chi$ | 17.6 $\alpha$ |

Variation is yield by season was outside the scope of our question and further compounded by differences in the number of picks between seasons. Therefore, yield data was not compared between seasons. Relative yield differences were observed in plants expressing *Bs2* compared to non-*Bs2* plants (Table 1). Only in Fall 2017 did *Bs2* plants statistically yield higher than non-*Bs2* plants. No yield difference was observed between plants expressing only *EFR* and WT plants in that season, but *Bs2* plants consistently had higher yields than *EFR* plants. While clear differences were observed for disease severity among genotypes in both 2018 seasons, differences in bacterial leaf spot symptoms did not translate to a statistically significant difference in fruit yield.

*Phyllosphere community structure differs by trial and Bs2 expression:* The richness of bacterial genera in each sample was influenced by the expression of *Bs2* and subsequent bacterial spot infection (Fig. 2A, Table S2). Plants expressing *Bs2* contained on average 54 more unique taxonomic identities than their non-*Bs2* counterparts. Communities associated with Bs2 plants had higher Shannon index values than their non-Bs2 counterparts (Fig. 2B).

**Figure 2.**
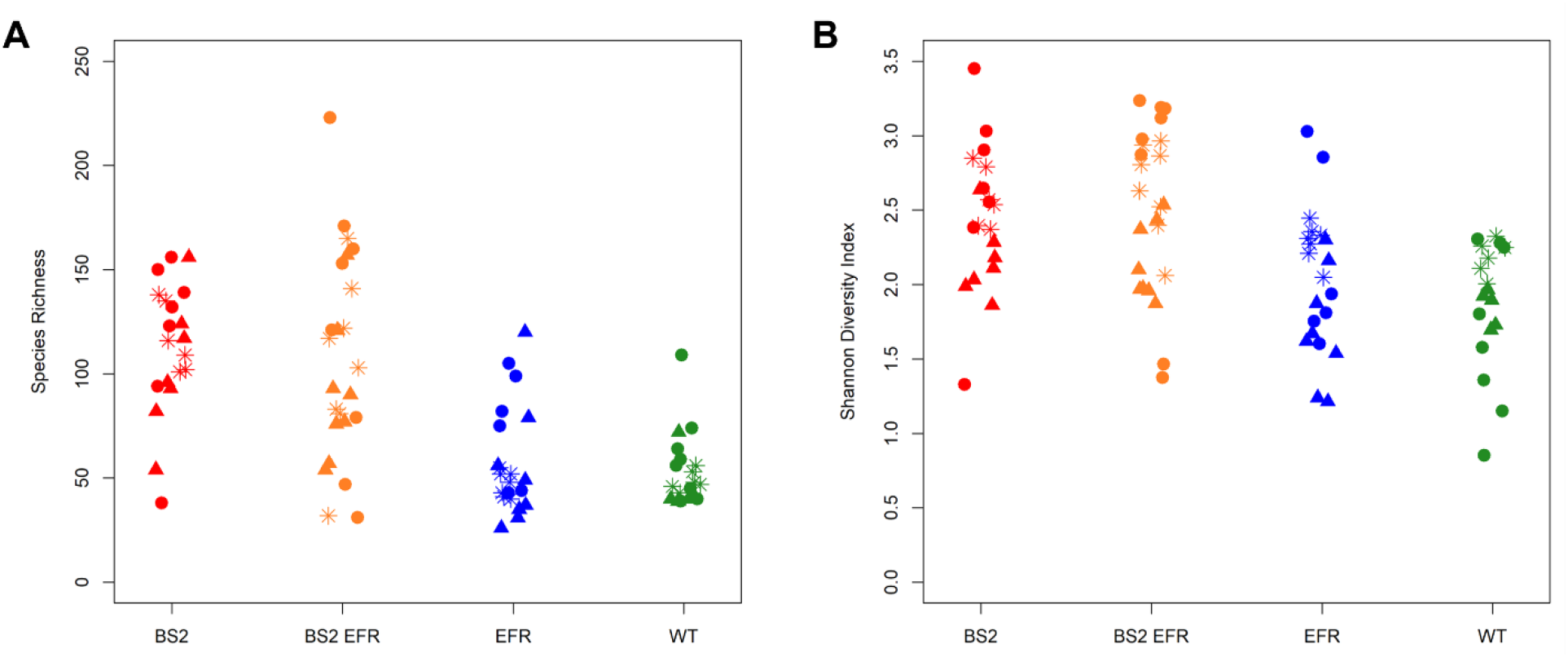
Disease decreases both species evenness and diversity. A one-way, analysis of variance (ANOVA) was used to assess alpha diversity, species richness (A) and Shannon Diversity (B). Color designates genotype (green, WT; blue, *EFR*; yellow *Bs2*/*EFR*; red, *Bs2*) and symbol sampling date (circle, fall 2017; asterisk, spring 2018; triangle, fall 2018).

A PERMANOVA of the Bray-Curtis distances revealed community structure significantly differed based on sampling date, *Bs2*, and an interaction between those two terms (p = 0.001). Visualization of Bray-Curtis distances using distance-based redundancy analysis (db-RDA) clustered *EFR* and WT plants together (Fig. 3). Combining the disease severity results, represented by the size of the symbol, with visualization of the db-RDA ordination (Fig. 3A) shows that similar communities on WT and *EFR* expressing plants were under high disease pressure (Fig. 3A). Plants expressing *Bs2* and low disease severity were separate from the WT and *EFR* only communities. Furthermore, *Bs2* communities from spring 2018 clustered separately from those from fall samplings in 2017 and 2018. The composition of bacterial communities of WT and *EFR* expressing plants differed between sampling dates. PERMANOVA of community composition with Xanthomonas ASVs removed from the dataset still displayed strong significant differences in community composition with respect to the BS2 gene and season, and no consistent effects of the EFR gene (Table S3).

**Figure 3.**
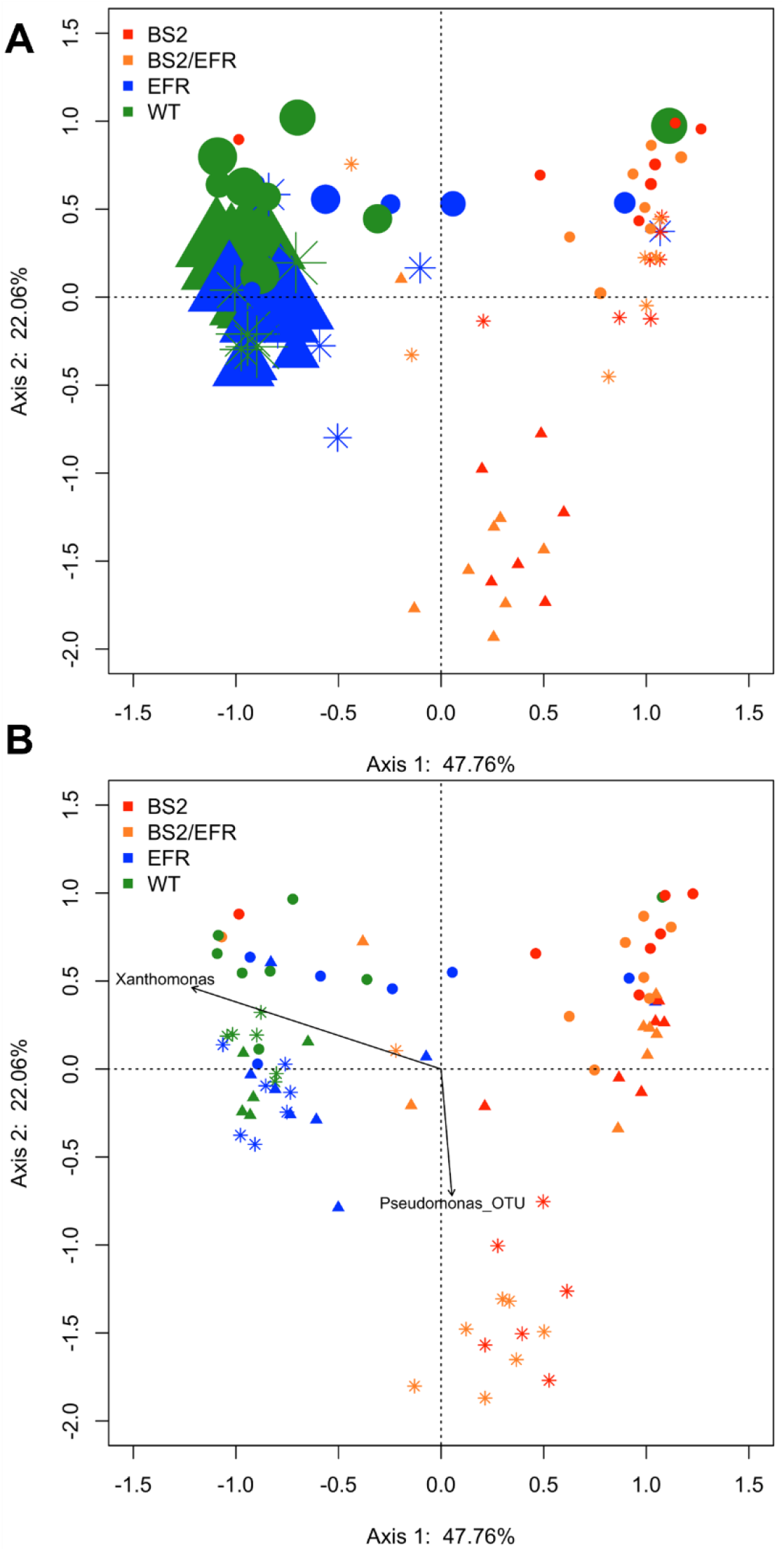
*Bs2* expression and season affect phyllosphere community structure. Pairwise comparisons in the overall structure of bacterial phyllosphere communities were made into a Bray-Curtis distance matrix and visualized using distance-based redundancy analysis. A) Disease severity results were combined with the Bray-Curtis distances to display disease pressure on bacterial communities of each genotype. B) Individual ASVs of interest, *Xanthomonas* and *Pseudomonas*, were fit onto the ordination as vectors. Color designates genotype (green, WT; blue, *EFR*; yellow *Bs2*/*EFR*; red, *Bs2*) and symbol sampling date (circle, fall 2017; asterisk, spring 2018; triangle, fall 2018).

Individual genera of interest, *Xanthomonas* and *Pseudomonas*, were fit onto the ordination as vectors (Fig. 3B). Just as high disease pressure was found in WT and *EFR* expressing plants, the ASV with an identical sequence to *X. perforans* used for inoculation was also associated with those bacterial communities. We chose to examine the ordination of the *Pseudomonas* ASV because symptoms of bacterial speck, caused by *P. syringae* pv. *tomato*, have been noted on *Bs2* expressing plants in spring plantings (G. Vallad, personal observation). The *Pseudomonas* ASV locates among the bacterial communities from *Bs2* expressing plants only in spring 2018. We found that 20 times more rain was recorded at the experimental site in May 2018 than either October 2017 or 2018, fall sampling months (Table S4). This increased rainfall likely influenced the increased bacterial spot severity recorded in the spring 2018 planting over the fall plantings (Fig. 1).

*Xanthomonas alters the phyllosphere in a way that only a small subset of taxa find conducive:* We used a mediation analysis to determine if the effect of the *Bs2* gene on community composition could be attributed in whole or part to its effect on disease severity. After accounting for the effects of seasons, the presence of the *Bs2* gene explained 23.7% of the variation in bacterial composition (Table S5). However, when disease severity was included first in the model, disease severity explained 25% of the variance in composition, and the *Bs2* gene explained only an additional 2.8% (Table S5). Very little of the variation in bacterial composition could be uniquely attributed to either the *Bs2* gene (2.8%) or disease severity (4%) – the vast majority of the effects of each of these two terms was shared with the other. As before, we repeated this analysis with a Xanthomonas filtered dataset. While the overall effect of the *Bs2* gene (and disease severity) was weaker (14.9 vs. 23.7% variation explained), the qualitative result was the same: most of the variation attributed to the *Bs2* gene could also be attributed to disease severity, with a small, but significant amount of variation attributed uniquely to the two terms (Table S4).

*Individual taxa differ in relative abundance based on the presence or absence of Bs2:* Because community structure was not altered by the expression of *EFR* as compared to WT, subsequent analysis of individual community members focuses on differences between *Bs2* and non-*Bs2* genotypes. In order to determine which taxa drive differences in community structure, Wilcoxon rank-sum tests were performed on all genera. Comparisons were made between *Bs2* genotypes (*Bs2, Bs2/EFR*) and non-*Bs2* (WT, *EFR*) genotypes. 604 genera were present in a sufficient number of samples to perform a Wilcox test. Of these 604 taxa, 119 differed in their rank sums between genotypes (FDR adjusted P value < 0.05). Taxa who differed by rank sum and had a greater relative abundance on *Bs2* genotypes were considered to be *Bs2* associated taxa (Fig. 4, checkered colors) whereas those with a higher abundance on non-*Bs2* plants were considered non-*Bs2* associated (Fig. 5, solid colors). A total of 106 taxa were *Bs2* associated; cumulatively, these taxa made up 51% of the community on *Bs2* plants and 13% on non-*Bs2* plants (Fig. 5). Excluding *Xanthomonas*, 12 of the 13 non-*Bs2* associated taxa made up 3.5% of *Bs2* communities and 16.5% of non-*Bs2* communities. Unassociated taxa, those with an FDR adjusted P value > 0.05, had similar distributions among genotypes, making up 36% and 32% of the *Bs2* and non-*Bs2* communities respectively.

**Figure 4.**
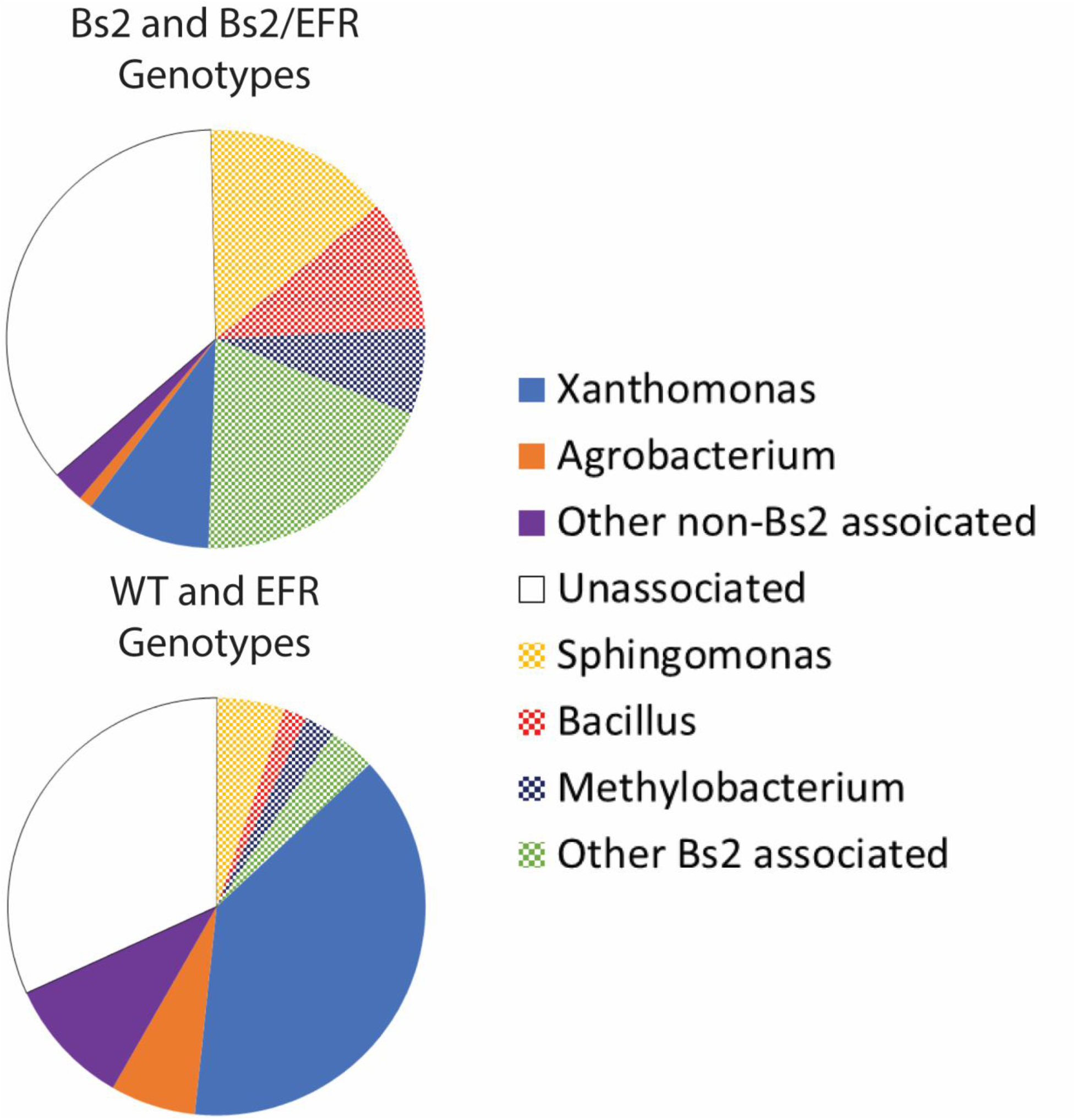
*Bs2* expression influenced individual taxa relative abundance. Wilcoxon rank-sum tests were performed on all genera and comparisons were made between *Bs2* and non-*Bs2* genotypes. Upper pie represents the proportion of taxa relative abundance found on *Bs2* and *Bs2*/*EFR* expressing plants while the lower pie is WT and *EFR* expressing plants. Taxa who differed by rank sum and had a greater relative abundance on *Bs2* genotypes labelled as *Bs2* associated (checkered colors) and those with a higher abundance on non-*Bs2* plants labelled non-*Bs2* associated (solid colors).

**Figure 5.**
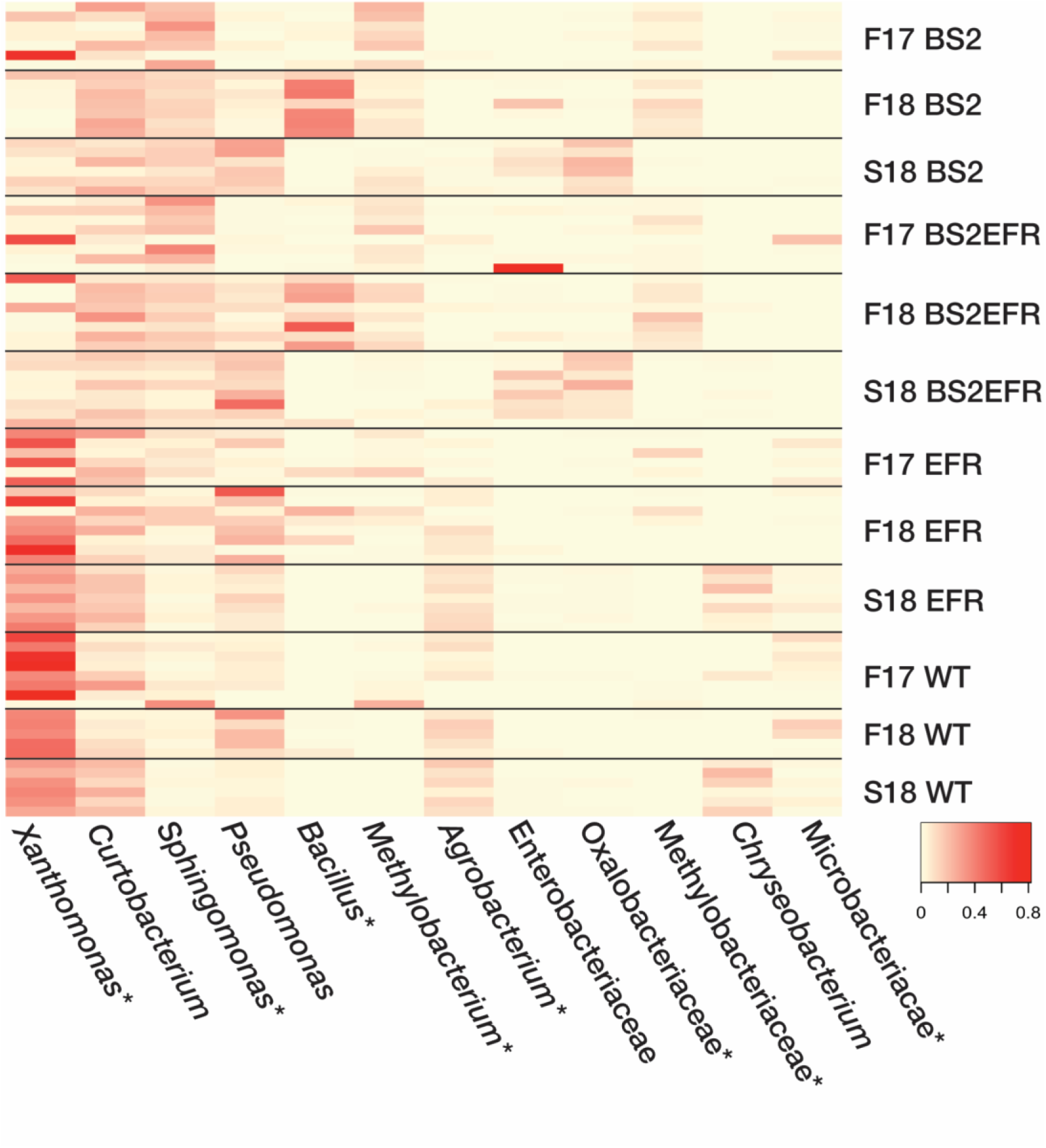
Twelve most abundant genera represented 85% of the overall phyllosphere bacterial community. Wilcoxon rank-sum tests were performed to determine general abundance; color intensity denotes relative abundance of each genera for each plot sampled. Asterisks represent taxa more closely associated with either *Bs2* or non-*Bs2* genotypes.

Across all genotypes, a small number of taxa made up the majority of the phyllosphere community. The twelve most relatively abundant genera represented 85% of the overall community (Fig. 5). *Xanthomonas* was found on plants of all genotypes and was the most relatively abundant member of the community from non-*Bs2* expressing plants. The proportion of *Xanthomonas* in the community was 4 times greater in non-*Bs2* plants. Of the 12 most relatively abundant taxa, 8 were either *Bs2* or non-*Bs2* associated (Fig. 5, denoted by asterisk). Common phyllosphere genera *Sphingomonas*, *Methylobacterium*, and *Pseudomonas* were among the most relatively abundant genera. Interestingly, *Pseudomonas* ranked as the third most relatively abundant genera across all genotypes and was not found associated with the presence or absence of *Bs2*, but made up 8.6% and 10.3% of *Bs2* and non-*Bs2* genotypes, respectively. Both *Sphigomonas* and *Methylobacterium* were among the most relatively abundant genera across all genotypes but were three times more relatively abundant on *Bs2* expressing plants. *Sphigomonas* and *Methylobacterium* relative abundance was negatively correlated with *Xanthomonas* relative abundance whereas *Agrobacterium* appears more abundant on non-*Bs2* plants along with *Xanthomonas*.

*Salmonella enterica identified as a member of the tomato phyllosphere bacterial community:* Three ASVs were identified as belonging to the genus *Salmonella.* In order to resolve the identity of these ASVs, sequences of closely related taxa were obtained from Genbank and the aligned sequences were used to construct a maximum likelihood phylogeny (Fig. S2). While not all *Salmonella* spp. form a single monophyletic clade, all *S. enterica* accessions formed a monophyletic clade along with the three ASVs putatively identified as such, suggesting an appropriate taxonomic identification. ASV 1 was the sequence most often identified as *S. enterica* whereas ASV 3 was only found in a single sample in spring 2018 from *Bs2/EFR* plants (Table S7). ASV 1 and ASV 2 were commonly found concurrently and seldomly in fall 2017 with the exception of three samples. In contrast, they were found in every sample in fall 2018 and every sample of *Bs2* and *Bs2/EFR* expressing plants in spring 2018.

## Discussion

Leaves support large populations of bacteria. The bacterial community of the tomato phyllosphere has previously been described as being similar to the airborne community (Ottesen et al. 2016) and stable, not prone to dramatic shifts from events such as irrigation (Telias et al. 2011). However, here we found that biotic stress, in the form of infection with *Xanthomonas* pathogens, resulted in dramatic restructuring of the phyllosphere community. The majority of individual bacterial genera were reduced in their relative abundance in diseased leaves, indicating that *Xanthomonas* infection alters the phyllosphere environment in ways that largely excludes non-pathogenic microbial inhabitants.

The presence of *Xanthomonas*, as well as the resulting bacterial spot disease, was the dominant factor influencing the bacterial community in this study. Due to the prevalence of *Xanthomonas spp.* in the environment as well as our experimental design, we found that yield, disease rating, and proportion of *Xanthomonas* in the metabarcoding were not affected by our experimental inoculation. Thus, we were unable to assess differences between bacterial communities of healthy and diseased WT or *EFR*-expressing plants. The overlap in terms of disease rating and *Bs2* in explaining variation in the alpha and beta diversity among samples suggests *Xanthomonas* pathogenesis leads to a less complex community dominated by the pathogen. Most of the variation in bacterial community composition that could be attributed to the *BS2* gene could also be attributed to variation in disease severity. The consensus of alpha and beta diversity mediation suggests bacterial spot is more directly responsible for phyllosphere community modification than is the subsequent resistance response. We found that only a small subset of phyllosphere taxa benefited from the disease state of the leaves. This is reflected in the depression in alpha diversity in diseased leaves and the finding that many more individual taxa were significantly more abundant on plants with the Bs2 resistance gene compared to the small number of non-*Bs2* associated taxa. Despite variation in healthy phyllosphere microbiomes from season to season, diseased microbiomes converged repeatedly into a similar community structure.

We found that while both *Bs2* and *EFR* expressing plants had significantly reduced bacterial spot, plants only expressing *EFR* had high levels of disease compared to *Bs2* expressing plants. Field-level resistance to bacterial spot has been demonstrated by plants expressing *Bs2*, *EFR*, or both (Horvath et al. 2012; Kunwar et al. 2018); however, the differences we observed in disease severity between *EFR* and *Bs2* expressing plants was similar to that reported earlier (Kunwar et al. 2018). In addition to resistance against bacterial spot, *EFR* expressing plants are resistant to bacterial wilt caused by *Ralstonia solanacearium* (Kunwar et al. 2018). Instead of recognition of a specific pathogen effector, *EFR* recognizes one of the most widely conserved and abundant proteins in bacteria (Kunze et al. 2004) and leads to downstream flux in ion concentrations, oxidative burst, and activation of mitogen-activated protein kinases (Jones and Dangl 2006). Somewhat surprisingly, the presence of the EFR transgene on its own did not result in substantial changes to the phyllosphere community structure (i.e. diversity or composition), perhaps because tomato plants expressing *EFR* do not appear to constitutively activate defense responses (Lacombe et al. 2010).

During infection, *Xanthomonas* fundamentally alters the phyllosphere. During disease development, *Xanthomonas* spreads from the point of entry and causes cell damage that extends beyond the site of infection (Ciardi et al. 2000). Cellular damage results in membrane instability and subsequent leakage of cellular constituents into the apoplast and onto the phylloplane. At the same time, leaves responsible for carbohydrate production through photosynthesis shift from carbohydrate sources to sinks (Garcia-Seco et al. 2017); infected leaves then receive carbohydrates from other source leaves instead of sending their own photosynthates away. These fundamental changes to the phyllosphere may in turn alter the distribution of nutrients across the infected leaf surface and the plant, in general. Nutrient oases are no longer restricted to glandular trichomes and cellular junctions, nutrient availability then follows Xanthomonas infection and phyllosphere colonization success may no longer be restricted to bacterial genera specialized in carbon scavenging and unique carbon utilization. Changes in carbon flow due to Xanthomonas pathogenesis may also trigger DAMP-mediated PTI. Therefore, copiotrophic microbes may need to subvert PTI in order to fully take advantage of liberated nutrients.

Changes to the available resource base in infected leaves may underlie the dramatic shifts in phyllosphere microbial composition. *Sphingomonas* and *Methylobacterium* are considered cosmopolitan members of the bacterial community due to their omnipresence in the phyllosphere across the plant kingdom. The dominance and commonality of Sphingomonas and Methylobacterium in phyllosphere bacterial communities across plant species has been attributed to these taxa’s capacity in carbon acquisition and/or utilization(Wilson and Lindow 1994; Delmotte et al. 2009). We found Sphingomonas and Methylobacterium among the most abundant genera on Bs2 expressing plants, which had little disease and smaller Xanthomonas relative abundance. Sphingomonas and Methylobacterium were not among the most abundant bacteria in the phyllosphere of non-Bs2 plants to the same extent that they were on healthy Bs2 plants. On the other hand, Agrobacterium relative abundance increased and was represented at a similar percentage to Sphingomonas and five times larger than Methylobacterium on the diseased plants without the Bs2 gene. This community structure shift suggests that on a diseased host, the capacity to use single carbon sugars no longer provided an overwhelming advantage.

Previous studies have examined how disease can alter the bacterial community of tomato, focusing on the root-associated communities, rhizoplane and rhizosphere, and effects of disease caused by *R. solanacearium* (Gu, Wei et al. 2016, Hu, Wei et al. 2016). In addition to the direct effects of disease on the plant, *R. solanacearium* infection alters root exudates which in turn directly affects the rhizosphere bacterial community.

Previous bacterial community studies of the tomato phyllosphere have also found an abundance of Xanthomonadaceae (Allard et al. 2018) and identified *Xanthomonas* as a core member of plants grown in Maryland, Virginia, and North Carolina, but not California (Ottesen et al. 2015, Ottesen et al. 2019). During the tomato growing season, weather conditions between California and the East Coast of the United States are distinct with low relative humidity and no rain events in California (https://www.wunderground.com/history). Environmental factors appear to strongly influence phyllosphere bacterial community structure.

Rain events have long been accepted to increase plant disease and food safety risks by enhanced dissemination of pathogens or creating a more favorable phyllosphere environment (Cevallos-Cevallos et al. 2012, Thompson et al. 2013). As the tomato grows and the phyllosphere physically moves up and away from the soil, the phyllosphere resembles the airborne community as opposed to the soil or rhizosphere community (Ottesen et al. 2013, Ottesen et al. 2016). A recent examination of airborne bacterial communities over a 7 year period found *Pseudomonas* as a core member and that consistent seasonal fluctuations of the airborne community can lead to bacterial introductions via rain events (Cáliz et al. 2018). We found that *Pseudomonas* was the third most abundant genera regardless of host genotype. A correlation unique to the tomato phyllosphere bacterial community was shown among members of *Pseudomonas* and *Methylobacterium* (Ottesen et al. 2016). We found the overall abundance of *Methylobacterium* associated with plants expressing *Bs2* but overall, we did not find *Pseudomonas* associated with host genotype when the fall samples were included. In spring 2018, we found a distinct community structure for plants possessing *Bs2* that were associated with a unique ASV, identified by 16S sequence as *P. syringae*, concurrently with a reported bacterial speck outbreak. A deluge of rainfall occurred during the month the spring samples were taken and the appearance of bacterial speck, caused by *Pseudomonas syringae* pv. tomato, was observed on *Bs2* expressing plants that season. While *Bs2* confers resistance against *Xanthomonas* infection, these plants are susceptible to *P. syringae* pv*. tomato* infection. In addition to a virulent pathogen and susceptible host, disease requires conducive environmental conditions such as optimal temperature and humidity (Horsfall and Dimond 1959) and microbial community (Hacquard et al. 2017). Optimal conditions for bacterial speck are high moisture, high relative humidity (> 80%) and cool temperature (17-23°C); the pathogen is inhibited during periods of high temperature (https://u.osu.edu/vegetablediseasefacts/tomato-diseases/bacterial-speck/advanced/). Plants resistant to *Xanthomonas* infection had a more diverse bacterial community and a second pathogen appeared to thrive under conducive environmental conditions and may have had its own impacts on phyllosphere bacterial communities.

Several ASVs identified as *Salmonella enterica* were found in samples from in fall 2018 and across all genotypes in spring 2018. We were surprised that the incidence of *S. enterica* was not correlated with plant disease as previous studies have identified plant disease as a food safety risk factor with larger populations of *S. enterica* found on diseased plants (Meng et al. 2013, Potnis al. 2014, Potnis et al. 2015). Given the capacity of *S. enterica* to thrive in the phyllosphere of *Xanthomonas-*infected tomato plants, we might have expected that genotypes without *Bs2* would be more likely hosts of *S. enterica.* However, in spring 2018 *Bs2* expressing plants were not disease free as bacterial speck was observed. Since we were unable to quantify the absolute abundance of *S. enterica* populations in the current study and only a single time point was sampled, the fate of the *S. enterica* populations among the host genotypes and whether plant disease affected them is unclear. It is also possible that our greater ability to detect *Salmonella* ASVs in the Bs2 genotypes may simply reflect the overwhelming abundance of *Xanthomonas* on the non-Bs2 genotypes, which inevitably reduces the relative abundance of the other microbial populations and can result in rarer taxa falling below our detection limit.

Previously, *S. enterica* was isolated from 2.2% of leafy greens, spinach, kale, and bok choy, sampled from farms in the mid-Atlantic region of the United States (Marine et al. 2015). Growing season (fall versus spring) was associated with *S. enterica* recovery, but not the farming system (organic or conventional) or region. The diversity in type of leafy greens samples, on-farm production practices, and geography suggests that a factor with widespread influence, such as climate, may have played a role in the importance of season and positive *S. enterica* leaves. More field-level research is needed to ascertain the importance of plant disease as a food safety risk factor, yet field research with human pathogens is restricted due to the human and environmental hazard.

In conclusion, we found effector-triggered immunity controlled the targeted plant disease while maintaining a diverse bacterial community. In fact, *Bs2* expressing plants consistently hosted *S. enterica* as a phyllosphere bacterial community member throughout two growing seasons and under conducive environmental conditions that permitted infection and disease by a second phytobacterial pathogen, *P. syringae* pv. *tomato*. Plants lacking this specific resistance suffered high pathogen loads and severe disease symptoms; these, in turn, led to a dramatic restructuring of the phyllosphere microbial community and a reduction in taxa adapted to life as leaf epiphytes.

## Notes

### Competing Interest Statement

The authors have declared no competing interest.

