## Supplemental Figures and Tables for "Narrow, but not broad, spectrum resistance and disease reshape phyllosphere bacterial communities"

**Supplemental Tables**

Table S1: ANOVA results for bacterial spot disease severity ratings.

| 3 Seasons Disease Severity Analysis | | | | | | |
|---|---|---|---|---|---|---|
| | Sum Sq | Mean Sq | NumDF | DenDR | F Value | Pr(>F) |
| BS2 | 24959 | 24959 | 1 | 81 | 200.922 | 2.20E-16 |
| EFR | 1704.2 | 1704.2 | 1 | 81 | 13.719 | 0.000386 |
| BS2:EFR | 1711 | 1711 | 1 | 81 | 13.773 | 0.000377 |
| Fall 2017 Disease Severity Analysis | | | | | | |
| | Sum Sq | Mean Sq | NumDF | DenDR | F Value | Pr(>F) |
| BS2 | 2083.63 | 2083.63 | 1 | 22.484 | 161.26 | 9.78E-12 |
| EFR | 289.49 | 289.49 | 1 | 22.484 | 22.404 | 9.55E-05 |
| BS2:EFR | 311.01 | 311.01 | 1 | 22.871 | 24.07 | 5.97E-05 |
| Fall 2018 Disease Severity Analysis | | | | | | |
| | Sum Sq | Mean Sq | NumDF | DenDR | F Value | Pr(>F) |
| BS2 | 3163.4 | 3163.4 | 1 | 24 | 264.632 | 1.83E-14 |
| EFR | 508.3 | 508.3 | 1 | 24 | 42.525 | 9.60E-07 |
| BS2:EFR | 495 | 495 | 1 | 24 | 41.412 | 1.18E-06 |
| Spring 2018 Disease Severity Analysis | | | | | | |
| | Sum Sq | Mean Sq | NumDF | DenDR | F Value | Pr(>F) |
| BS2 | 21097.4 | 21097.4 | 1 | 20.183 | 3.54E+02 | 2.84E-14 |
| EFR | 654.2 | 654.2 | 1 | 20.327 | 1.10E+01 | 0.0034 |
| BS2:EFR | 703.2 | 703.2 | 1 | 20.183 | 1.18E+01 | 0.002584 |

Table S2. P-values for alpha diversity analyses.

| | Effects of genotype across all sampling dates | | Effects of *Bs2* expression for individual seasons | | |
|---|---|---|---|---|---|
| | *Bs2* | *EFR* | Fall 17 | Spring 18 | Fall 18 |
| Species Richness | 5.8e-10 | 0.997 | 2.25e-03 | 1.18e-06 | 1.03e-03 |
| Shannon Diversity | 1.07e-06 | 0.295 | 5.13e-03 | 8.35e-05 | 8.01e-04 |

Table S3. perMANOVA results of phyllosphere bacterial community composition excluding *Xanthomonas.*

| | Df | Sums of Sqs | Mean Sps | F. Model | R2 | Pr(>F) | |
|---|---|---|---|---|---|---|---|
| Season | 2 | 3.1811 | 1.59057 | 15.0964 | 0.20376 | 0.001 | *** |
| BS2 | 1 | 2.3327 | 2.33274 | 22.1406 | 0.14942 | 0.001 | *** |
| EFR | 1 | 0.0653 | 0.0653 | 0.6198 | 0.00418 | 0.75 | |
| Season:BS2 | 2 | 1.9227 | 0.96136 | 9.1245 | 0.12316 | 0.001 | *** |
| Season:EFR | 2 | 0.1658 | 0.08289 | 0.7867 | 0.01062 | 0.666 | |
| BS2:EFR | 1 | 0.1626 | 0.16259 | 1.5432 | 0.01041 | 0.129 | |
| Season:BS2 | 2 | 0.1956 | 0.0978 | 0.9282 | 0.01253 | 0.508 | |
| Residuals | 72 | 7.586 | 0.10536 | 0.48591 | | | |
| Total | 83 | 15.6118 | 1 | | | | |
| Fall 2017 | | | | | | | |
| BS2 | 1 | 0.7584 | 0.75838 | 4.3351 | 0.14234 | 0.006 | ** |
| EFR | 1 | 0.0803 | 0.08031 | 0.4591 | 0.01507 | 0.889 | |
| BS2:EFR | 1 | 0.1156 | 0.11563 | 0.6609 | 0.0217 | 0.657 | |
| Residuals | 25 | 4.3735 | 0.17494 | 0.82088 | | | |
| Total | 28 | 5.3278 | 1 | | | | |

| | | | | | | | |
|---|---|---|---|---|---|---|---|
| Spring 2018 | | | | | | | |
| BS2 | 1 | 2.0744 | 2.07443 | 30.1523 | 0.53921 | 0.001 | *** |
| EFR | 1 | 0.0794 | 0.07944 | 1.1546 | 0.02065 | 0.268 | |
| BS2:EFR | 1 | 0.1109 | 0.11093 | 1.6123 | 0.02883 | 0.183 | |
| Residuals | 23 | 1.5824 | 0.0688 | 0.41131 | | | |
| Total | 26 | 3.8472 | 1 | | | | |
| Fall 2018 | | | | | | | |
| BS2 | 1 | 1.4253 | 1.42534 | 20.9855 | 0.4378 | 0.001 | *** |
| EFR | 1 | 0.0686 | 0.06864 | 1.0106 | 0.02108 | 0.34 | |
| BS2:EFR | 1 | 0.1316 | 0.13164 | 1.9381 | 0.04043 | 0.121 | |
| Residuals | 24 | 1.6301 | 0.06792 | 0.50069 | | | |
| Total | 27 | 3.2557 | 1 | | | | |

Table S4. Monthly rainfall (mm) measured at 2 m height at the Gulf Coast Research Center in Balm, FL. https://fawn.ifas.ufl.edu/data/

| | 2017 | 2018 |
|---|---|---|
| March | 17.78 | 29.21 |
| April | 69.85 | 85.34 |
| May | 44.96 | **328.68** |
| June | 261.62 | 175.01 |
| July | 247.14 | 277.37 |
| August | 199.14 | 201.42 |
| September | 226.57 | 116.33 |
| October | 14.73 | 13.97 |
| November | 3.302 | 55.37 |

Table S5. Effects of BS2 gene versus bacterial spot disease severity on bacterial community composition.

| | Effect | Df | SS | PseudoF | $R^2$ | P |
|---|---|---|---|---|---|---|
| Model 1 | Season | 2 | 2.7316 | 11.96 | 0.16876 | 0.001 |
| | Disease Severity | 1 | 4.0417 | 35.392 | 0.24971 | 0.001 |
| | BS2 | 1 | 0.45 | 3.941 | 0.0278 | 0.011 |
| | EFR | 1 | 0.055 | 0.482 | 0.0034 | 0.807 |
| | Residuals | 78 | 8.9076 | | 0.55033 | |
| | Total | 83 | 16.186 | | 1 | |
| Model 2 | Season | 2 | 2.7316 | 11.96 | 0.16876 | 0.001 |
| | BS2 | 1 | 0.651 | 5.701 | 0.04022 | 0.002 |
| | Disease Severity | 1 | 3.8407 | 33.632 | 0.23729 | 0.001 |
| | EFR | 1 | 0.055 | 0.482 | 0.0034 | 0.807 |
| | Residuals | 78 | 8.9076 | | 0.55033 | |
| | Total | 83 | 16.186 | | 1 | |
| Non-Xanthomonas community | | | | | | |
| Model 1 | Season | 2 | 3.1811 | 13.4919 | 0.20376 | 0.001 |
| | Disease Severity | 1 | 2.6317 | 22.3233 | 0.16857 | 0.001 |
| | BS2 | 1 | 0.5248 | 4.4513 | 0.03361 | 0.001 |
| | EFR | 1 | 0.0787 | 0.6678 | 0.00504 | 0.676 |
| | Residuals | 78 | 9.1955 | | 0.58901 | |
| | Total | 83 | 15.6118 | | 1 | |
| Model 2 | Season | 2 | 3.1811 | 13.4919 | 0.20376 | 0.001 |
| | BS2 | 1 | 2.3327 | 19.7873 | 0.14942 | 0.001 |

| | | | | | | |
|---|---|---|---|---|---|---|
| | Disease Severity | 1 | 0.8237 | 6.9873 | 0.05276 | 0.001 |
| | EFR | 1 | 0.0787 | 0.6678 | 0.00504 | 0.708 |
| | Residuals | 78 | 9.1955 | | 0.58901 | |
| | Total | 83 | 15.611 | | 1 | |

Table S6. Accessions of full-length 16S rRNA genes from Greengenes used to construct a maximum likelihood phylogeny. http://greengenes.secondgenome.com

| Accession | Taxonomy |
|---|---|
| 4475936 | *Salmonella enterica* serovar Typhi |
| 4350788 | *Salmonella ente*rica serovar Schwarzengrund |
| 4306121 | *Salmonella enterica* serovar Enteritidis |
| 558281 | *Serratia marcescens* |
| 291478 | *Klebsiella pneumonia* |
| 737912 | *Citrobacter freundii* |
| 687940 | *Enterobacter arachidis* |
| 4359731 | *Pantoea agglomerans* |
| 656881 | *Escherichia coli* |
| 114510 | *Escherichia coli* |
| 775089 | *Dikeya chrysanthemi* |
| 274754 | *Yersinia kristensenii* |

Table S7. Percentage of specific sequences from the bacterial community identified as *Salmonella enterica*.

| Sample | ASV 1 | ASV 2 | ASV 3 |
|---|---|---|---|
| F17 Bs2 | 0 | 0 | 0 |
| F17 Bs2 | 0 | 0 | 0 |
| F17 Bs2 | 0 | 0 | 0 |
| F17 Bs2 | 0 | 0 | 0 |

| | | | |
|---|---|---|---|
| F17 Bs2 | 0 | 0 | 0 |
| F17 Bs2 | 0 | 0 | 0 |
| F17 Bs2 | 0 | 0 | 0 |
| F17 Bs2/EFR | 0 | 0 | 0 |
| F17 Bs2/EFR | 0 | 0 | 0 |
| F17 Bs2/EFR | 0 | 0 | 0 |
| F17 Bs2/EFR | 0 | 0 | 0 |
| F17 Bs2/EFR | 0 | 0 | 0 |
| F17 Bs2/EFR | 0 | 0 | 0 |
| F17 Bs2/EFR | 0 | 0 | 0 |
| F17 Bs2/EFR | 0 | 0 | 0 |
| F17 EFR | 0 | 0 | 0 |
| F17 EFR | 0 | 0 | 0 |
| F17 EFR | 0 | 0 | 0 |
| F17 EFR | 0.0019 | 0.0029 | 0 |
| F17 EFR | 0.0049 | 0.0016 | 0 |
| F17 EFR | 0 | 0 | 0 |
| F17 EFR | 0 | 0 | 0 |
| F17 WT | 0 | 0 | 0 |
| F17 WT | 0 | 0 | 0 |
| F17 WT | 0 | 0 | 0 |
| F17 WT | 0 | 0 | 0 |
| F17 WT | 0.0007 | 0 | 0 |
| F17 WT | 0 | 0 | 0 |
| F17 WT | 0 | 0 | 0 |
| F17 WT | 0 | 0 | 0 |
| S18 Bs2 | 0.0082 | 0.0027 | 0 |
| S18 Bs2 | 0.0103 | 0.0024 | 0 |

| | | | |
|---|---|---|---|
| S18 Bs2 | 0.0095 | 0.0021 | 0 |
| S18 Bs2 | 0.0089 | 0.0024 | 0 |
| S18 Bs2 | 0.0138 | 0.008 | 0 |
| S18 Bs2 | 0.0054 | 0.0016 | 0 |
| S18 Bs2/EFR | 0.0005 | 0.0001 | 0 |
| S18 Bs2/EFR | 0.0009 | 0.0003 | 0 |
| S18 Bs2/EFR | 0.0019 | 0.0003 | 0 |
| S18 Bs2/EFR | 0.0001 | 0.0001 | 0 |
| S18 Bs2/EFR | 0.0047 | 0.0009 | 0 |
| S18 Bs2/EFR | 0.0004 | 0.0002 | 0 |
| S18 Bs2/EFR | 0.0015 | 0.0012 | 0 |
| S18 Bs2/EFR | 0.1594 | 0.0531 | 0.0045 |
| S18 EFR | 0 | 0 | 0 |
| S18 EFR | 0 | 0 | 0 |
| S18 EFR | 0 | 0 | 0 |
| S18 EFR | 0.0002 | 0.0001 | 0 |
| S18 EFR | 0.0004 | 0.0001 | 0 |
| S18 EFR | 0 | 0 | 0 |
| S18 WT | 0 | 0 | 0 |
| S18 WT | 0 | 0 | 0 |
| S18 WT | 0 | 0 | 0 |
| S18 WT | 0 | 0.0001 | 0 |
| S18 WT | 0 | 0 | 0 |
| S18 WT | 0 | 0 | 0 |
| F18 Bs2 | 0.0122 | 0.004 | 0 |
| F18 Bs2 | 0.0177 | 0.0063 | 0 |
| F18 Bs2 | 0.0142 | 0.0049 | 0 |
| F18 Bs2 | 0.0018 | 0.0006 | 0 |

| | | | |
|---|---|---|---|
| F18 Bs2 | 0.0041 | 0.0048 | 0 |
| F18 Bs2 | 0.0033 | 0.0013 | 0 |
| F18 Bs2 | 0.0027 | 0.002 | 0 |
| F18 Bs2/EFR | 0.017 | 0.0006 | 0 |
| F18 Bs2/EFR | 0.0034 | 0.0019 | 0 |
| F18 Bs2/EFR | 0.0005 | 0 | 0 |
| F18 Bs2/EFR | 0.0079 | 0.0036 | 0 |
| F18 Bs2/EFR | 0.0105 | 0.0018 | 0 |
| F18 Bs2/EFR | 0.0084 | 0.0043 | 0 |
| F18 Bs2/EFR | 0.0041 | 0.0015 | 0 |
| F18 Bs2/EFR | 0.0026 | 0.0012 | 0 |
| F18 EFR | 0.0002 | 0 | 0 |
| F18 EFR | 0.0003 | 0 | 0 |
| F18 EFR | 0.0023 | 0 | 0 |
| F18 EFR | 0.0012 | 0.0003 | 0 |
| F18 EFR | 0.0003 | 0 | 0 |
| F18 EFR | 0.0002 | 0.0001 | 0 |
| F18 EFR | 0.0006 | 0 | 0 |
| F18 EFR | 0.0002 | 0 | 0 |
| F18 WT | 0.0001 | 0.0002 | 0 |
| F18 WT | 00.002 | 0 | 0 |
| F18 WT | 0 | 0.0006 | 0 |
| F18 WT | 0.0001 | 0.0001 | 0 |
| F18 WT | 0.0019 | 0.0004 | 0 |

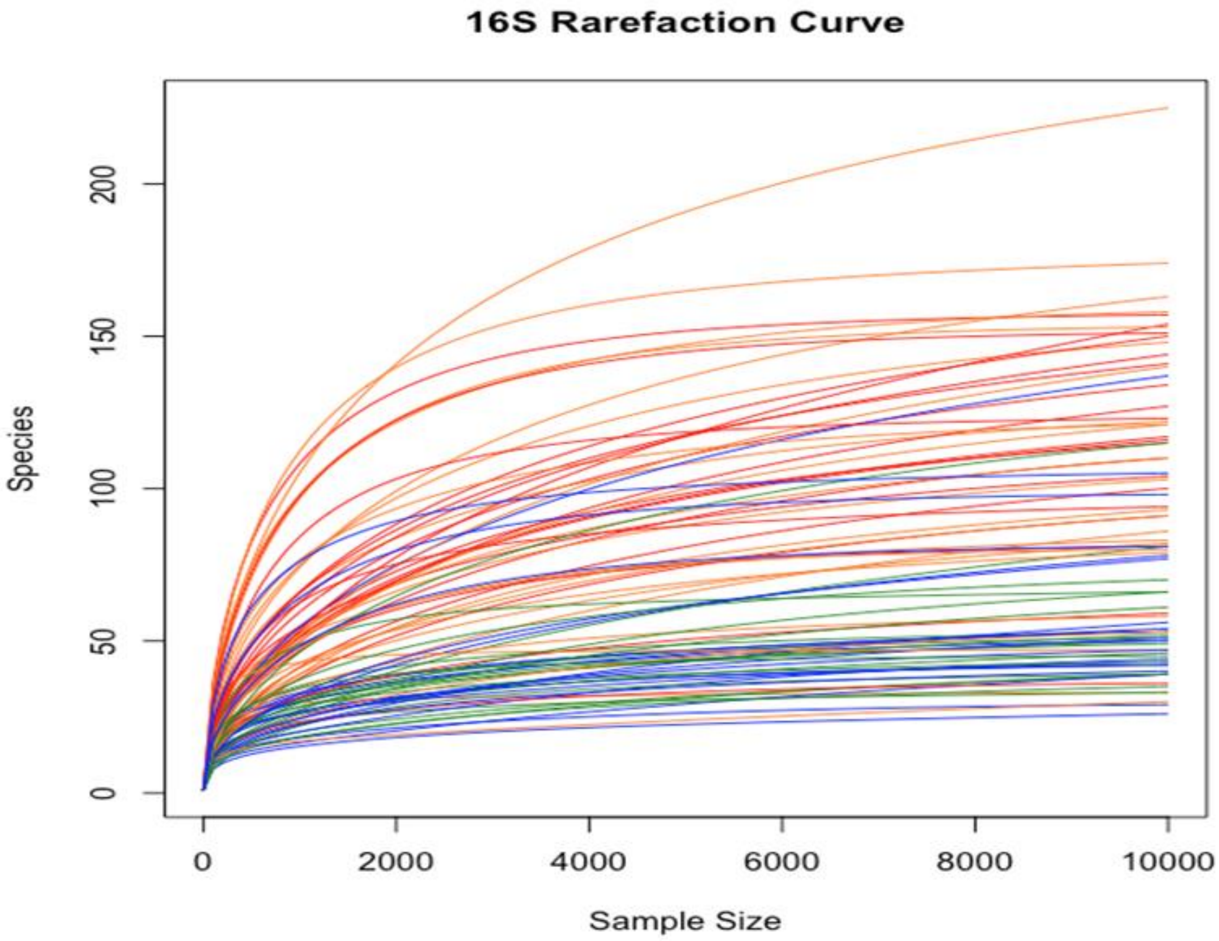

Figure S1: Rarefaction curves of the number of genera identified per sample

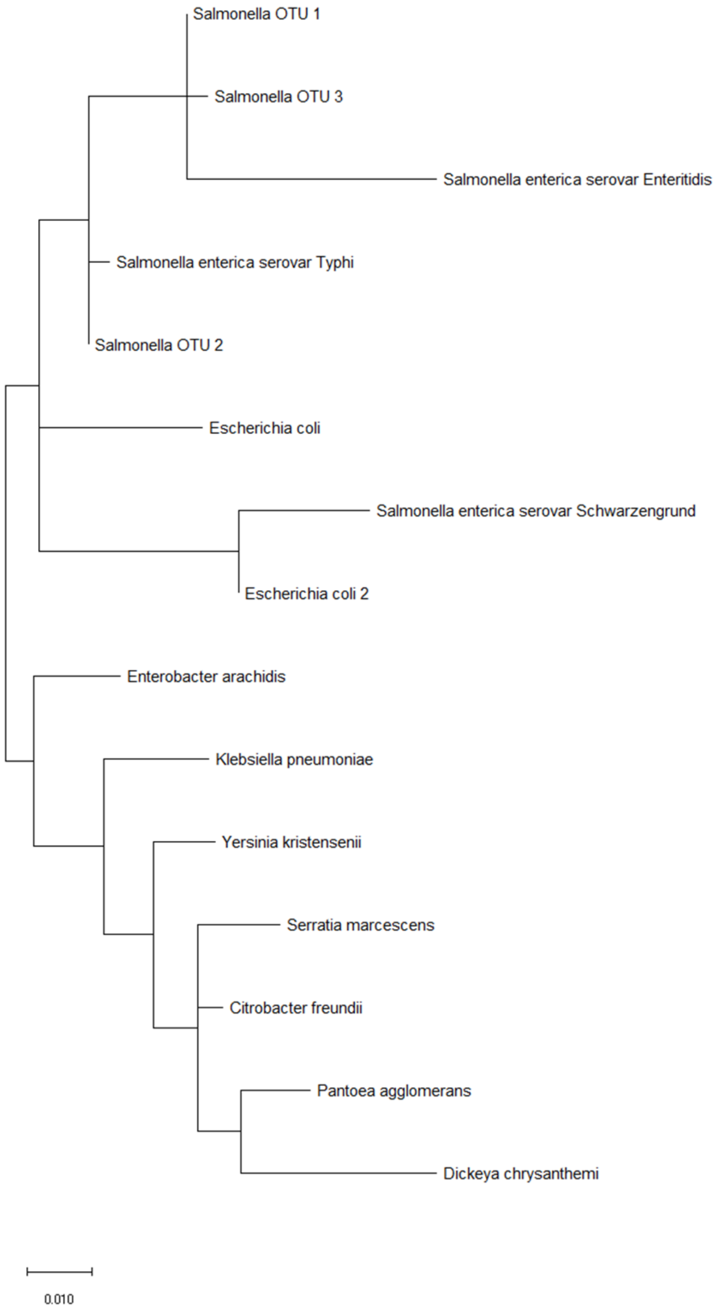

Figure S2: Maximum likelihood phylogeny of *Salmonella* enterica sequences and three ASVs identified as belonging to the genus *Salmonella*.
